# The OS-Prey (Omnibus Study of Prey) database: A compilation of diet records for birds of prey

**DOI:** 10.64898/2026.03.28.714998

**Authors:** Stella F Uiterwaal, Frank A La Sorte, Kyle E Coblentz, John P DeLong

## Abstract

**Motivation:** The diet composition of a predator is a direct reflection of its role in a food web, resulting from interactions with prey species. Raptors (including hawks, owls, and falcons) are ubiquitous predators with diverse diets, yet there is no comprehensive database of raptor diet composition. We present a database of over 3500 raw raptor diet records, compiled from more than 1000 studies and representing 173 raptor species from across the world. Our dataset complements existing qualitative summaries of species diets by compiling thousands of quantitative diet “samples” over time and space to present diet data at a uniquely fine resolution.

**Main types of variable contained:** The database comprises published records of raptor diets from pellets, prey remains, direct or photographic observations, prey DNA, and raptor gut or gullet contents. For each diet, we present the taxonomic identity and amounts of consumed prey. We additionally present various metadata for each diet such as location, habitat, and season.

**Spatial location and grain:** The study incorporates diet records collected worldwide, with each record assigned geographic coordinates corresponding to the location where the diet information was obtained.

**Time period and grain:** The database includes diet records from 1893 to 2025. We report a year for each diet record.

**Major taxa and level of measurement:** We recorded raptor diet at the species level, including raptors from three orders: Strigiformes, Falconiformes and Accipitriformes excluding vultures. Most prey are identified to species, but prey taxonomic level varies depending on the extent to which they could be identified.

**Software format:** Diet records and metadata are provided in two files with comma-separated value (.csv) format.

## Introduction

Diets - the identity and quantity of consumed prey - are a key component of predators’ ecology. By capturing trophic links between predators and prey, diets describe food web structure and reveal how species interact with each other and their shared environment (DeLong, 2021; McCann, 2011). Because diets are the mechanism by which energy and biomass flow through food webs, diets are also linked to populations dynamics and individual fitness (Rosenblatt and Schmitz, 2016; Yodzis and Innes, 1992). Predation thus also steers evolutionary trajectories, including the development of morphological traits and behavioral strategies that underlie diets (Caro, 2005; Lima and Dill, 1990; Slagsvold et al., 2010). The study of predator diets is therefore integral to our understanding of ecology.

Raptors (hawks, falcons, and owls) are conspicuous and influential predators in communities worldwide, making the study of raptor diets integral to understanding how these communities function. Their diets reflect patterns of prey availability, habitat quality, and species interactions that may otherwise be difficult to detect. Raptors’ ecological importance is amplified by their highly mobile nature: many raptors migrate long distances or use expansive home ranges, connecting food webs across ecosystems (Bildstein, 2006; DeLong et al., 2024). Raptor diets also provide a window into environmental health. Because their feeding ecology exposes them to bioaccumulated toxins, raptor diets are intimately linked to major conservation threats such as DDT-induced eggshell thinning, lead poisoning from ammunition, and secondary rodenticide poisoning (Katzner et al., 2024; Nakayama et al., 2019; Olsen et al., 1993).

Because of this, researchers have been interested in what raptors eat for decades, making raptors the focus of extensive dietary data collection efforts. This research has provided insight into countless ecological questions related to foraging mechanisms, migration strategies, prey populations, and environmental effects (Bednarz et al., 1985; Bourbour et al., 2019; Teta et al., 2012; Whalen et al., 2024) and contributed to the conservation and management of both raptor and prey species (Seamans and Gutierrez, 1999). Yet, despite the wealth of raptor diet literature, there is not a centralized resource for accessing this data. Individual diet studies have generated invaluable insights, but their scope is often limited in taxonomic, geographic, or temporal coverage. Some compilations exist, but these focus on the diets of certain species (e.g., Obuch, 2011) or of raptors in a specific community (e.g., Korschgen and Stuart, 1972). We lack a global compilation of raptor diet research that would allow us to use the existing body of research to its fullest potential. To meet this need, we here present the Omnibus Study of Prey (OS-Prey) database, a standardized compilation of raptor diet records worldwide.

## Methods

### Diet record search

We used a multi-pronged approach to search for diet records in an attempt to maximize the number of publications we encountered (van den Burg and Kaiser, 2026). We searched Google Scholar and Web of Science, as well as ornithology and regional zoological journals using the keywords “diet”, “raptor”, and “pellets”. We additionally searched for specific well-studied species, such as “barn owl”. Last, we looked through the references of raptor diet papers to find additional sources. We included records that included a diet (i.e., a list of prey taxa and quantities) for a known raptor species (i.e., in the orders Falconiformes, Strigiformes, and Accipitriformes, but excluding vultures). We excluded records where the raptor species was unknown, prey numbers were not quantified, prey taxa were extremely broad (e.g., none of the prey identified to species), or prey taxa were systematically excluded from the record. Although most publications were written in English, we also obtained raptor diet data from articles written in Spanish, French, Italian, Portuguese, and German. We included records published through 2025.

### Diet extraction

We recorded prey taxa and quantities from the original records. We preferentially recorded quantities as number of prey but occasionally report only relative frequencies if total prey number was not reported. We included records at the highest level of detail reported in the study. For example, if a publication reported both an overall diet as well as seasonal diet, we recorded the seasonal data. We additionally recorded the taxonomic resolution of each prey type as well as the class, the order, and a common name if applicable. If prey were known to be eggs, we recorded this as well.

### Diet metadata

We recorded a source for each diet, including the study (authors and year) and publication. When a source reported a diet originally published elsewhere, we attempted to locate the original source and used the secondary version only if the original source could not be located. In this case, we noted both sources. We recorded the raptor species, genus, family, order, and common name in English.

We additionally categorized the method of diet collection as reported in the source. “Pellets” indicated diet data collected from regurgitated pellets. “Prey remains” denotes identifications from carcasses, bones, feathers, or fur not obtained from pellets. “Direct observations” refer to hunting, consumption, or prey deliveries noted by a human observer, while “Photos/videos” refer to prey documented by camera. “Stomach/gullet contents” refer to diet data obtained from material found in raptors’ stomachs or gullets, such as from birds killed by vehicle collisions. “DNA” indicates diet data obtained from DNA such as from swabs of prey remains on the talons or beak. If data was collected only from pellets, we also recorded the total number of pellets (if available) and the mean prey per pellet. We either calculated prey per pellet by dividing the number of prey by the number of pellets or reported the prey per pellet as reported in the source.

We recorded the country and continent where the data was collected, as well as the latitude and longitude. If coordinates were provided in the source, we used this location. Otherwise, we estimated coordinates from maps provided in the source or using location cues (e.g., nearest town) reported in the source. We also reported the habitat associated with the raptor diet using study site descriptions to determine which landscape categories a diet originated from: riparian/lake, woodland/forest/rainforest, orchard/plantation forest, pasture/grazed land, prairie/grass/meadow, scrub/shrub/heath, urban, agriculture, wetland/marsh, desert, tundra, coastal/beach/island. Because study site descriptions in the literature are not systematic, this classification is intended to provide qualitative ecological context for the diets.

We additionally reported temporal data for each diet. If diet data was from the breeding season (e.g., prey deliveries to nestlings), non-breeding season (e.g., wintering owls), or year round, we recorded this. We additionally recorded the quarter of the year in which the data was collected (Dec-Feb, Mar-May, Jun-Aug, Sept-Nov) to provide seasonal context for the data. Lastly, we recorded a year for each diet, taking the mean if data was collected across multiple years. We note that this year is distinct from the publication year.

## Results

The resulting database consists of 3,538 raptor diet records across 173 raptor species (Fig. 1A). While some raptors are very well represented in OS-Prey (*Tyto alba, Strix aluco*, and *Asio otus* each had more >100 diet records) there are only a handful of diets for most species (Fig. 2A).

**Figure 1.**
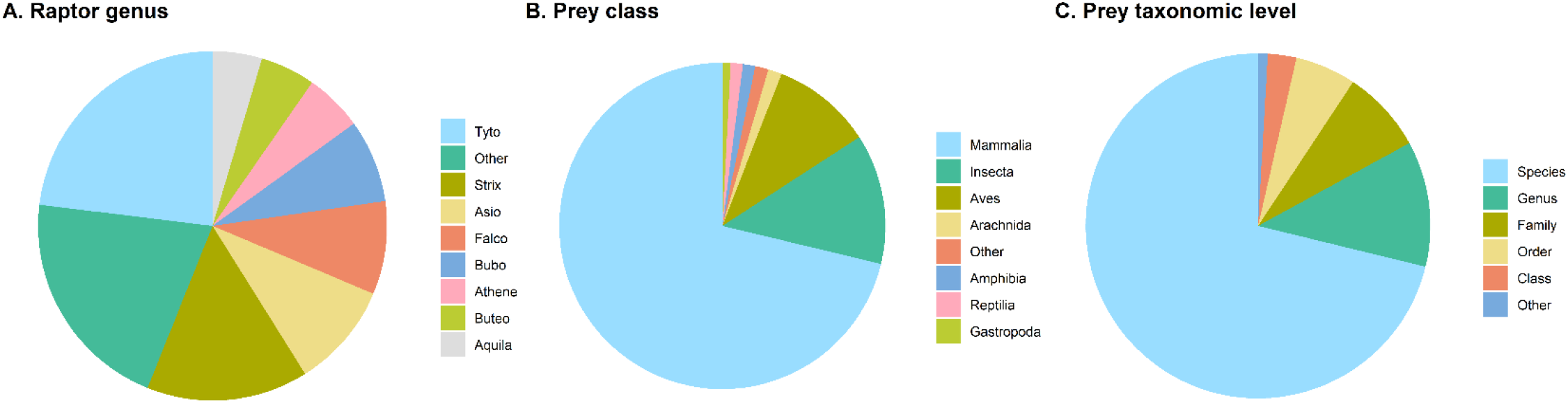
A) Proportion of diet records in OS-Prey by raptor genus. B) Proportion of prey classes across all raptor prey reported in OS-Prey. C) Finest taxonomic resolution of all raptor prey reported in OS-Prey.

**Figure 2.**
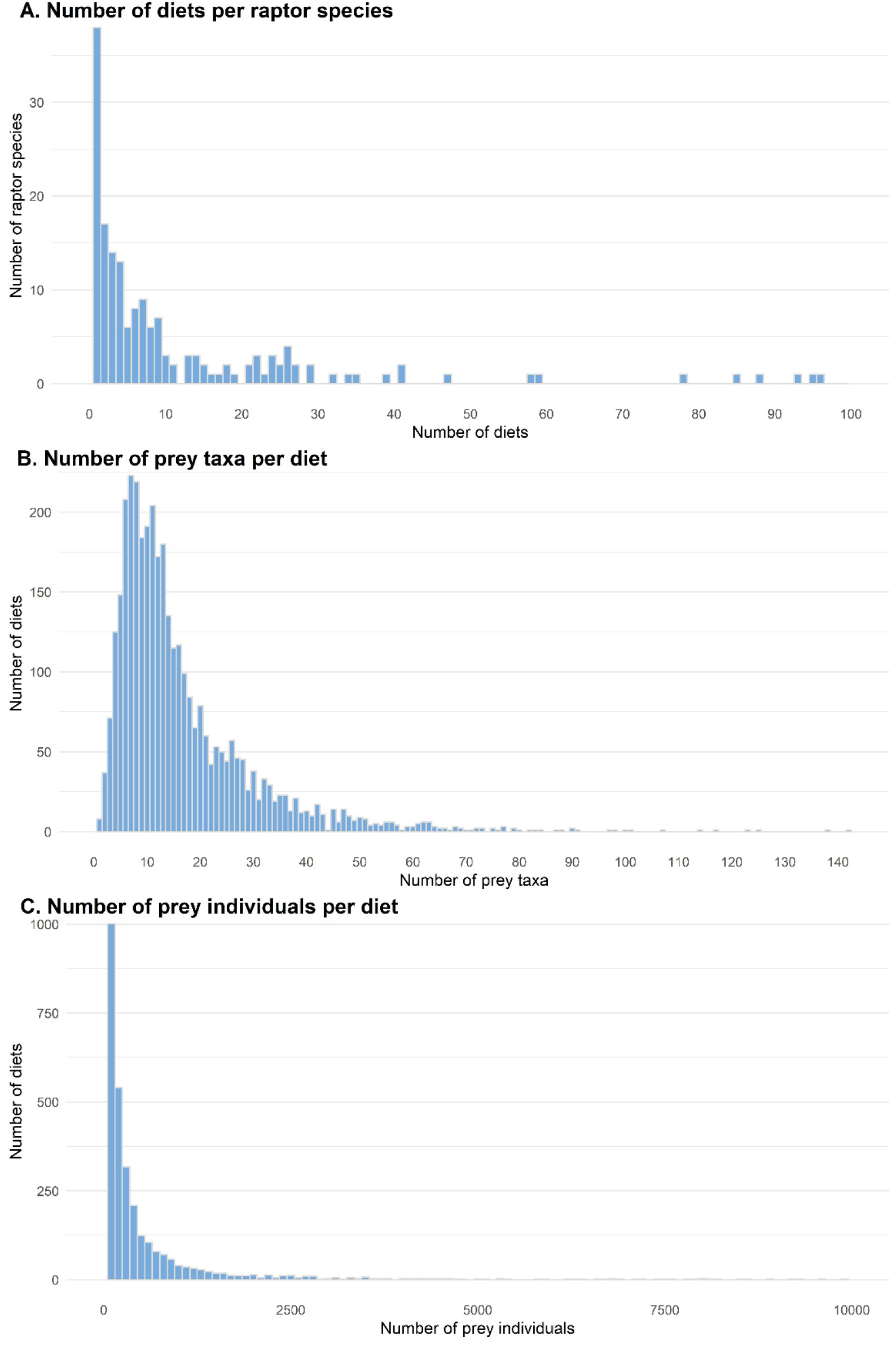
A) Number of studies for each raptor species in OS-Prey. Three species with many studies are not shown: *Tyto alba*, 799 studies; *Strix aluco*, 429 studies; *Asio otus*, 251 studies. B) Number of prey taxa reported per study in OS-Prey. C) Number of prey individuals per diet in OS-Prey. 23 diets records each containing more than 10,000 prey individuals are not shown.

A total of 2,555,630 prey individuals belonging to 5,206 prey taxa are documented in OS-Prey. Prey are mostly mammals with insects and birds also commonly reported (Fig. 1B). Most prey types could be identified to species, but some prey are identified to a coarser taxonomic resolution (Fig. 1C). Diet records contain a median of 12 taxa (Fig. 2B) and 178 prey individuals (Fig. 2C).

The database is available in two comma-separated value (.csv) files from the Knowledge Network for Biocomplexity (Uiterwaal et al., 2026). RaptorDiets.csv provides data on the study, raptor species, prey taxa, and number or frequency of prey. RaptorDiets_Metadata.csv provides data on the study, raptor taxa, location, habitat, season, year, and diet collection method.

## Discussion

The OS-Prey database is the largest compilation of raptor diet records to date. Its scale is possible because researchers around the world have invested extraordinary effort in documenting what raptors consume, with the sheer number of diet records and prey items illustrating how extensively raptor foraging ecology has been studied. OS-Prey underscores the substantial investment by the global raptor research community that made a dataset of this magnitude possible. At the same time, the uneven distribution of records across taxa highlights how much effort has been concentrated on certain species, with many species represented by just a handful of diet records.

As new studies continue to be published on raptor diets, we intend to provide future updated versions of OS-Prey by including diets published after 2025. We encourage authors of such studies to contact us for inclusion of their datasets. We also intend to use these updates to supplement OS-Prey with any additional datasets we encounter. We attempted to identify as many published raptor diets as possible by using a variety of discovery methods. However, as with any literature search, there are undoubtedly articles that we missed and we also encourage authors whose studies were omitted to contact us. Providing updated versions of OS-Prey will allow us to include these additional datasets.

The OS-Prey database underscores the central ecological role raptors play as predators across the globe. OS-Prey reveals extensive variation in the number of taxa contained in raptor diets, with many diets including a dozen or more taxa. Across studies, these diets show both the strong reliance of many raptors on mammals and as well as the remarkable breadth of prey taxa consumed by the raptor guild as a whole, including insects, birds, fish, and reptiles. This diversity highlights raptors as important consumers whose foraging patterns can be used to explore broader patterns of biodiversity, prey availability, and environmental change (DeLong et al., 2024).

By bringing together thousands of diet records, OS-Prey creates an opportunity to build on our current understanding of predation by enabling researchers to explore high-levels questions such as determinants of diet breadth or the phylogeny of diet composition. OS-Prey also opens the door to more broad-scale conservation applications: researchers can use it to assess ecosystem health or explore how raptors respond to shifting landscapes. Because raptors diets are samplers of the species they forage on (Heisler et al., 2016; Terry, 2010; Viteri et al., 2022), OS-Prey could even be used to assess communities of rodents or other common prey across the world. Together, these possibilities position OS-Prey as a foundation for many future studies that links predator ecology, prey dynamics, and ecosystem change, and we encourage researchers to take full advantage of its potential.

## Data availability statement

The OS-Prey database is available from KNB at knb.ecoinformatics.org/view/urn%3Auuid%3A3e99f6df-c95a-47b1-999d-cdb7c12e5edd

